# Lower performance of *Toxoplasma*-infected, Rh-negative subjects in the weight holding and hand-grip tests

**DOI:** 10.1101/264093

**Authors:** Jaroslav Flegr, Blanka Šebánková, Lenka Příplatová, Veronika Chvátalová, Šárka Kaňková

## Abstract

*Toxoplasma*, a protozoan parasite of cats, infects many species of intermediate and paratenic hosts, including about one-third of humans worldwide. After a short phase of acute infection, the tissue cysts containing slowly dividing bradyzoites are formed in various organs and toxoplasmosis proceeds spontaneously in its latent form. In immunocompetent subjects, latent toxoplasmosis was considered asymptomatic. However, dozens of studies performed on animals and humans in the past twenty years have shown that it is accompanied by a broad spectrum of specific behavioural, physiological and even morphological changes. In human hosts, the changes often go in the opposite direction in men and women, and are mostly weaker or non-existent in Rh-positive subjects. Here, we searched for the indices of lower endurance of the infected subjects by examining the performance of nearly five hundred university students tested for toxoplasmosis and Rh phenotype in two tests, a weight holding test and a grip test. The results confirmed the existence of a negative association of latent toxoplasmosis with the performance of students, especially Rh-negative men, in these tests. Surprisingly, but in an accordance with some already published data, *Toxoplasma*-infected, Rh-positive subjects expressed a higher, rather than lower, performance in our endurance tests. Therefore, the results only partly support the hypothesis for the lower endurance of *Toxoplasma* infected subjects as the performance of Rh-positive subjects (representing majority of population) correlated positively with the *Toxoplasma* infection.

## Introduction

About one third of the world population are infected with the protozoan parasite *Toxoplasma gondii* (Tenter *et al.,* 2000). In immunocompetent subjects, after a short phase of acute toxoplasmosis promoted by tachyzoites, the disease spontaneously proceeds into its latent phase, which is characterized by the presence of tissue cysts with a slowly dividing form of the parasite, bradyzoites, in various organs and anamnestic IgG antibodies in blood (Pappas *et al.,* 2009). Latent toxoplasmosis has been considered asymptomatic for a long time, however, during the past 20 years, about one hundred papers have been published showing that *Toxoplasma* seropositive and seronegative subjects differ in many personality traits (Flegr & Hrdý, 1994; Flegr *et al.,* 2003), performance in certain psychomotor tests (Havlíček *et al.,* 2001; Pearce *et al.,* 2013), cognitive tests (Dickerson *et al.,* 2014; Ferreira *et al.,* 2013; Flegr *et al.,* 2012; Pearce *et al.,* 2014) and also in the incidence and form of many diseases and disorders (Flegr & Escudero, 2016; Flegr *et al.,* 2014). In some tests, however, *Toxoplasma-infected* subjects score better than *Toxoplasma-free* individuals (Flegr *et al.,* 2013; Stock *et al.,* 2014). Very often, these differences deepen with time since the infection (Flegr *et al.,* 2000; Flegr *et al.*, 1996; Havlíček *et al.*, 2001) and the analogical changes were nearly always reported to occur in experimentally infected laboratory animals (Hodková *et al.,* 2007; Vyas, 2015; Vyas & Sapolsky, 2010; Webster, 2001). This suggests that the infection is most probably the cause of the behavioral changes, rather than that the special combination of phenotypic traits is the cause of the infection. It has been suggested that these observed behavioral differences are either the product of the manipulation activity of the parasite aimed to increase the probability of transmission of *Toxoplasma* from an intermediate to a definitive host by predation, or potentially as side-effects of the pathological changes accompanying the infection. Very often, personality differences associated with toxoplasmosis differ in scope or even direction between men and women. The stress-coping hypothesis (Lindová *et al.,* 2010; Lindová *et al.*, 2006) explains the opposite reactions of men and women to *Toxoplasma* infection by the fact that men and women cope in an opposite way to mild chronic stress. In contrast to stressed men, who use more individualistic and antisocial (for example aggressive) forms of coping (Carver *et al.,* 1989; Hobfoll *et al.,* 1994), stressed women more often seek and provide social support (Stone and Neale 1984, Rosario et al. 1988, Carver et al. 1989), join with others (Hobfoll *et al.,* 1994), and verbalize towards others or ones’ self (Tamres *et al.,* 2002).

Similarly, *Toxoplasma* infection is associated with different phenotypical outputs in Rh-positive and Rh-negative subjects. The Rh-positive subjects, especially Rh-positive heterozygotes, seem to be fully or partly protected against many of the effects of toxoplasmosis, such as personality changes (Flegr *et al.,* 2010), decreased psychomotor performance (Flegr *et al.,* 2008c; Novotná *et al.,* 2008), and impaired physical and mental health (Šebánková & Flegr, 2017). The strongest effect of latent toxoplasmosis reported until now, the excessive weight gain of *Toxoplasma* infected women in the 16^th^ week of their pregnancies (4.12 kg vs 2.44 kg, N=152, 25 Toxoplasma-infected), was observed only in Rh-negative mothers (Kaňková *et al.,* 2010).

One of the earliest reported effects of latent toxoplasmosis in humans capable of enhancing the transmission of the parasite by predation in infected animals was the phenomenon of lower endurance of infected individuals. In three independent questionnaire surveys, infected subjects, especially men, responded affirmatively to six of ten statements of the Toxo-92 questionnaire significantly more often, including the following item: “When I am attacked, physically or otherwise, or when I should fight for something important, I stop fighting at one moment. It is not a result of a rational decision not to fight, as in fact I know that I should continue fighting and I would like to do so, but my own subconsciousness betrays me and I loss the will to fight back”(Flegr, 2010). It was suggested that the reason of this premature surrender could be the lack of endurance in *Toxoplasma-infected* subjects. However, no experimental data or other empirical evidence, possibly except an increased rate of suicides or suicide-attempts in the infected subjects (Arling *et al.,* 2009; Ling *et al.,* 2011; Okusaga *et al.,* 2011; Yagmur *et al.,* 2010; Zhang *et al.,* 2012), have thus far been published.

The aim of present study was to search for empirical evidence for a decreased will to fight in the *Toxoplasma-infected* subjects. In double-blind experiments, we measured the endurance of *Toxoplasma-infected* and *Toxoplasma-free* university students using two tests – a weight-holding tests and a hand-grip test. As the effects of toxoplasmosis very often differ in men and women and in Rh-positive and Rh-negative subjects, we included the variables sex and Rh phenotype into our statistical models.

## Material and methods

### Population

The population consisted mostly of students of biology of the Faculty of Science, Charles University, who were recruited during undergraduate and graduate level courses of Evolutionary Biology and Methodology of Science between 2010-2015. Usually, more than 70 % of students attending these coursed accepted the invitation to participate in projects studying the manipulation activity of the parasite *Toxoplasma.* They were offered the opportunity to enrol in such experiments during a course entitled Methods of Evolutionary and Experimental Psychology, and compensated for their participation with one credit toward their assessment. The project, including the method of recruitment of participants and obtaining informed consent, was approved by the Institutional Review Board of the Faculty of Science, Charles University (No. 2014/21).

Registered probands signed an informed consent, completed several questionnaires including an anamnestic questionnaire, and provided 3 ml of blood for testing for the presence of anamnestic anti-*Toxoplasma* antibodies and for testing of Rh phenotype. Experiments were separated to three sessions and took about ten hours altogether. Probands participated in all the sessions within several weeks. Both of the tests of performance were completed in the first session, which took place once a week and always started at 9:00 am. After finishing the whole battery of tests, they were informed about results of their serological tests for toxoplasmosis and Rh factor and rewarded with 400 CZK for their participation in the experiment.

First, probands got an email invitation to attend a session ahead of time and chose a date according to their preferences. They were asked to fulfil some requirements: to sleep sufficiently during the night before testing; to refrain from physical exertion for several days beforehand; to cancel their attendance in case of an illness, physical or mental problems or convalescence; to not drink coffee, tea, alcohol, and other energy drinks the day of the experiment. They were, however, offered a snack to replenish their energy during the long experimental session.

### Grip test

Hand grip dynamometry is one of the screening procedures measuring fitness in a normal population. Hand grip strength can predict vitality, the nutritional state of an individual, as well as possible functional bodily failure (Bohannon, 2001; Mafi *et al.,* 2012). A Collin dynamometer, a type of closed steel spring dynamometer, was used for testing grip strength. Hand grip strength was measured in kg.

The test was performed first in a battery of experiments. Each proband was requested to stand upright, with arms kept freely along their trunk, and to look straight ahead. They took the gauge by their dominant arm and the experimenter explained how they should assumed a fixed position in their hand. The proband was asked to exert the maximal practicable effort on the grip without moving their trunk and shoulders. After the performance, the experimenter recorded the measurement and asked the proband to repeat the sequence two more times with the same hand, and then three times with their nondominant hand. After the test was over, the experimenter informed the proband of his results and told him which of his or her hands was stronger. This part of the experiment was executed individually in the same room with other probands, but in seclusion.

### Weight-holding test

A weight-holding performance test was prepared as a secondary experiment to be implemented in the same session as the grip test. It was placed at the end of a day-long experimental session, preceded by an intelligence and memory test, and several questionnaires distributed in paper and electronic forms. The probands were accommodated with two breaks and a snack in between the parts of the session. Each proband was requested to stand upright, not to lean over or backward, and to look straight ahead with experimenters outside his or her visual field. The proband was fitted with a hinged weight (5 kg) and asked to grasp it with both hands, to tense their arms in a levelled position with their shoulders, and with the backs of their hands up. The timer (measurement in seconds) was started by an experimenter the moment the appropriate position was engaged. The test was finished by the proband or by the experimenter when the proband was no longer capable of stabilizing their arms in a levelled position with shoulders. The probands were informed of their results after completing the test.

### Testing for toxoplasmosis and Rh phenotype

Testing for toxoplasmosis was performed at the National Reference Laboratory for Toxoplasmosis, National Institute of Public Health, Prague. The complement-fixation test (CFT), which determines the overall levels of IgM and IgG antibodies of particular specificity, and Enzyme-Linked Immunosorbent Assays (ELISA) (IgG ELISA: SEVAC, Prague) were used to detect the *T. gondii* infection status of the subjects. ELISA assay cut-point values were established using positive and negative standards according to the manufacturer’s instructions. In CFT test, the titre of antibodies against *Toxoplasma* in sera was measured in dilutions between 1:4 and 1:1024. The subjects with CFT titres between 1:8 and 1:128 were considered *Toxoplasma* infected (the subjects with higher titres, who were suspected to be in the acute phase of the infection, were excluded from the study). Only subjects with clearly positive or negative results of CFT and IgG ELISA tests were diagnosed as *Toxoplasma-infected* or *Toxoplasma-free,* whilst subjects with different results on these tests, or ambiguous results, were retested or excluded from the study. A standard agglutination method was used for Rh factor examination. A constant amount of anti-D serum (human monoclonal antiD reagent; SeracloneH, Immucor Gamma Inc.) was added to a drop of blood on a white glass plate. Red cells of Rh-positive subjects were agglutinated within 2–5 min.

### Statistics

Statistic analyses were performed with program Statistica v. 10.1. The effect of *Toxoplasma* seropositivity and Rh on participants’ performance in the weight-holding test was measured with ANCOVA (modul General Linear Models), with the independent binary variables *Toxoplasma* seropositivity (toxo), Rh phenotype (Rh), and sex of a subject (sex), the covariate age of a subject in time of testing (age), and toxo-Rh, toxo-sex, Rh-sex, and Rh-toxo-sex interactions as the independent variables. The effect of toxoplasmosis and Rh on participants’ performance in the hand-grip test was measured with repeated measures ANCOVA, with the performance of particular subjects in six trials (3 dominant hand trials and 3-nondominant hand trials) as the repeated measures; the independent variables were toxo, Rh, sex, and two nested factors: hand (dominant/nondominant) and trial (1^st^, 2^nd^, and 3^rd^ trial for each hand). The model also included all interactions of toxo, Rh, sex, hand, and trial. In the follow-up analyses, the same tests (without the factor sex and without all interactions with sex) were performed separately for men and women.

All raw data are available at Figshare https://figshare.com/s/692c3a7d08c3df860c9e

### Terminological notes

For the sake of clarity, we abbreviated “*Toxoplasma* seropositivity” to “toxoplasmosis” in the Discussion and to “toxo” in the description of our statistical models. Also, the statistical relations between (formally) dependent and (formally) independent variables are called “effects,” despite the fact that the real causal relation between these variables can be different or even non-existent (as reminded in the Discussion).

## Results

### Weight-holding test

This test was performed by 289 women, 59 (20.4%) *Toxoplasma-seropositive* and 176 men, 29 (16.5%) Toxoplasma-seropositive. The seroprevalence of toxoplasmosis did not differ between women and men (p = 0.339, Fisher exact test). Similarly, frequency of Rh-negative subjects, 21.1% (61) in women and 19.3% (34) in men, did not differ between men and women or between *Toxoplasma-seropositive* and Toxoplasma-seronegative male or female subjects (Chi^2^ = 1.31, df = 4, p = 0.860, all partial p values > 0.30). The age of *Toxoplasma-seropositive* subjects was higher than age of seronegative subjects (women: 24.6, SD 6.5 vs 22.6, SD =3.30, t_287_ = −3.39, p = 0.001, men: 26.4, SD 9.00 vs 23.9, SD =4.56, t_174_ = −2.19, p = 0.029). The effect of Toxoplasma-seropositivity and Rh phenotype in the weight-holding test was measured with GLM with time of holding as the dependent variable and toxo, Rh, age, sex, toxo-Rh, toxo-sex, Rh-sex, toxo-Rh-Sex as independent variables. Age and especially sex had very significant effects on participants’ performance in the test (age: eta^2^ = 0.023, p = 0.0004; sex: eta^2^ = 0.20, p < 0.0005). The effect of toxo depended on the Rh phenotype of subjects (toxo-Rh interaction: eta^2^ = 0.011, p = 0.026). Toxoplasma-seropositive, Rh-negative subjects expressed lower performance and *Toxoplasma-seropositive* Rh-positive subjects expressed higher performance than their Toxoplasma-seronegative peers, see the Fig. 1. Separate GLM analyses performed for men and women showed that the effect was significant neither for men, nor for women (p values > 0.10).

**Fig. 1.**
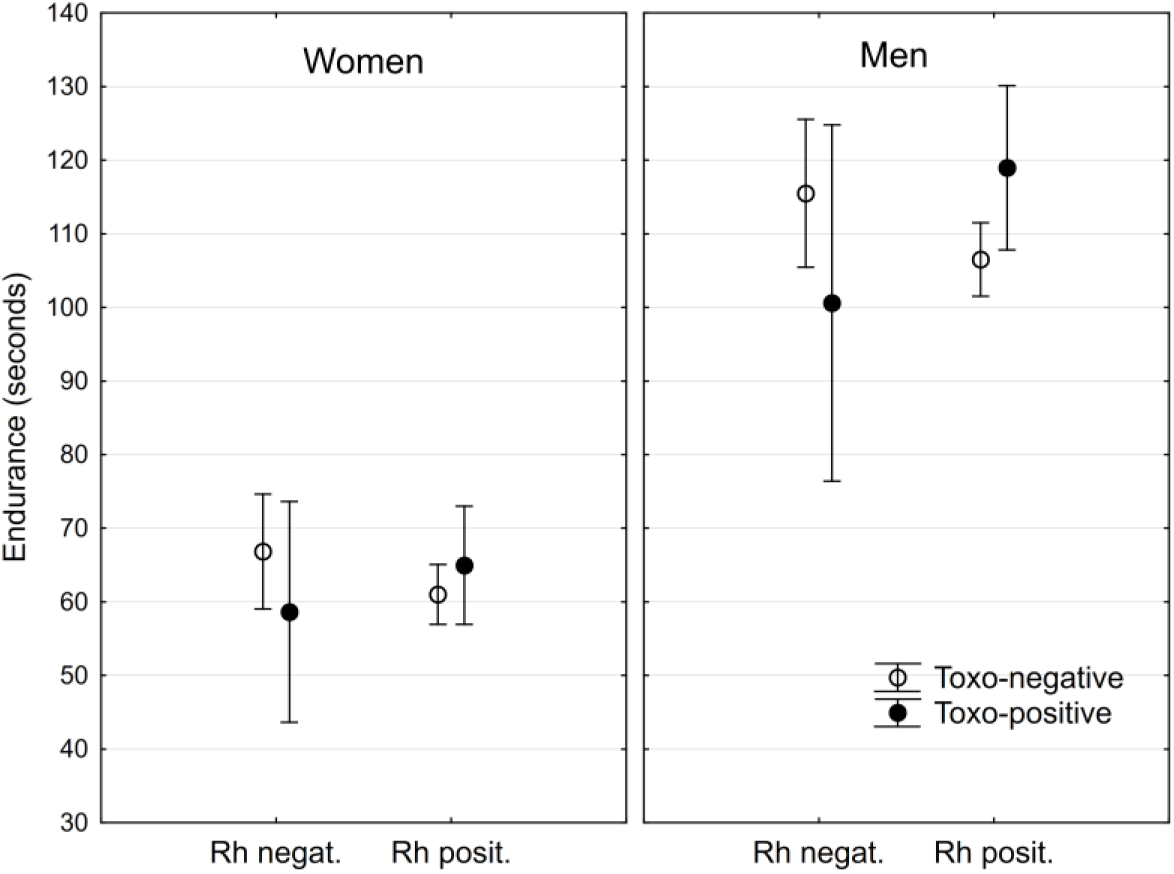
Effects of toxoplasmosis and Rh phenotype on performance in the weight holding test. Points show arithmetic means of endurance computed for covariate at its mean and spreads denote 95% confidence intervals.

### Hand-grip test

The hand-grip test was performed by 343 women, 66 (19.2 %) *Toxoplasma-seropositive* and 207 men, 35 (16.9 %) Toxoplasma-seropositive. The seroprevalence of toxoplasmosis did not differ between women and men (p = 0.570, Fisher exact test). Similarly, the frequency of Rh-negative subjects, 21.9% in women and 18.8% in men, did not differ between men and women or between *Toxoplasma*-seropositive and Toxoplasma-seronegative male or female subjects (Chi^2^ = 1.48, df = 4, p = 0.830, partial p values > 0.11). The age of *Toxoplasma-seropositive* subjects was higher than age of seronegative subjects (women: 24.2, SD 6.3 vs 22.48, SD = 3.09, t_345_ = −3.24, p = 0.001, men: 25.8, SD 8.38 vs 23.7, SD = 4.45, t_206_ = −2.12, p = 0.035). The effects of Toxoplasma-seropositivity and Rh phenotype in the hand-grip test were measured with repeated measures GLM with the performance in three dominant hand-trials and in three non-dominant hand-trials as nested repeated measures, and toxo, Rh, sex, hand, trial, age, and all interactions of toxo, Rh, sex, hand, and trial as independent variables. Table 1 shows the results of this analysis and analyses performed separately for men and women. Fig. 2 shows that the effect of trial-Rh-toxo is stronger in men than in women and that the toxoplasmosis had negative effects on the performance of the Rh-negative subjects. Separate analyses performed for men and women showed that the effects were significant for men, but not for women, see Table 1.

**Table 1.** Effects of toxoplasmosis, Rh phenotype, hand and trial on performance in the hand-grip test. Table shows the results (p values and partial η^2^) of repeated measures analyses of variances performed for all subjects and also separately for women and men. Each participant had three trials with their dominant (either right or left) and three trials with their non-dominant hand. Significant results are printed in bold; p values < 0.00005 are coded as 0.000.

|  | All |  | Women |  | Men |  |
| --- | --- | --- | --- | --- | --- | --- |
| | p | $\eta^2$ | p | $\eta^2$ | p | $\eta^2$ |
| Age | **0.000** | **0.027** | **0.003** | **0.025** | **0.015** | **0.029** |
| Sex | **0.000** | **0.298** |  |  |  |  |
| Rh | 0.721 | 0.000 | 0.560 | 0.001 | 0.897 | 0.000 |
| Toxo | 0.355 | 0.002 | 0.441 | 0.002 | 0.562 | 0.002 |
| Sex-Rh | 0.933 | 0.000 |  |  |  |  |
| Sex-Toxo | 0.795 | 0.000 |  |  |  |  |
| Rh-Toxo | 0.175 | 0.003 | 0.364 | 0.002 | 0.351 | 0.004 |
| Sex-Rh-Toxo | 0.487 | 0.001 |  |  |  |  |
| Hand | **0.000** | **0.037** | **0.000** | **0.042** | **0.007** | **0.035** |
| Hand-Age | 0.535 | 0.001 | 0.158 | 0.006 | 0.888 | 0.000 |
| Hand-Sex | **0.000** | **0.034** |  |  |  |  |
| Hand-Rh | 0.064 | 0.006 | 0.179 | 0.005 | **0.025** | **0.025** |
| Hand-Toxo | 0.334 | 0.002 | 0.461 | 0.002 | 0.247 | 0.007 |
| Hand-Sex-Rh | **0.003** | **0.016** |  |  |  |  |
| Hand-Sex-Toxo | 0.093 | 0.005 |  |  |  |  |
| Hand-Rh-Toxo | 0.202 | 0.003 | 0.266 | 0.004 | 0.115 | 0.012 |
| Hand-Sex-Rh-Toxo | **0.031** | **0.009** |  |  |  |  |
| Trial | **0.000** | **0.015** | **0.008** | **0.014** | **0.028** | **0.018** |
| Trial-Age | 0.429 | 0.002 | 0.428 | 0.003 | 0.782 | 0.001 |
| Trial-Sex | **0.032** | **0.006** |  |  |  |  |
| Trial-Rh | **0.001** | **0.012** | 0.109 | 0.007 | **0.027** | **0.018** |
| Trial-Toxo | **0.029** | **0.007** | 0.783 | 0.001 | **0.024** | **0.018** |
| Trial-Sex-Rh | 0.157 | 0.003 |  |  |  |  |
| Trial-Sex-Toxo | **0.012** | **0.008** |  |  |  |  |
| Trial-Rh-Toxo | **0.001** | **0.014** | 0.199 | 0.005 | **0.012** | **0.022** |
| Trial-Sex-Rh-Toxo | 0.088 | 0.004 |  |  |  |  |
| Hand-Trial | 0.092 | 0.004 | 0.276 | 0.004 | 0.314 | 0.006 |
| Hand-Trial-Age | 0.194 | 0.003 | 0.246 | 0.004 | 0.623 | 0.002 |
| Hand-Trial-Sex | 0.074 | 0.005 |  |  |  |  |
| Hand-Trial-Rh | 0.051 | 0.005 | 0.850 | 0.000 | 0.060 | 0.014 |
| Hand-Trial-Toxo | 0.502 | 0.001 | 0.282 | 0.004 | 0.178 | 0.009 |
| Hand-Trial-Sex-Rh | 0.075 | 0.005 |  |  |  |  |
| Hand-Trial-Sex-Toxo | **0.034** | **0.006** |  |  |  |  |
| Hand-Trial-Rh-Toxo | 0.062 | 0.005 | 0.583 | 0.002 | **0.024** | **0.018** |
| Hand-Trial-Sex-Rh-Toxo | **0.009** | **0.009** |  |  |  |  |

**Fig. 2.**
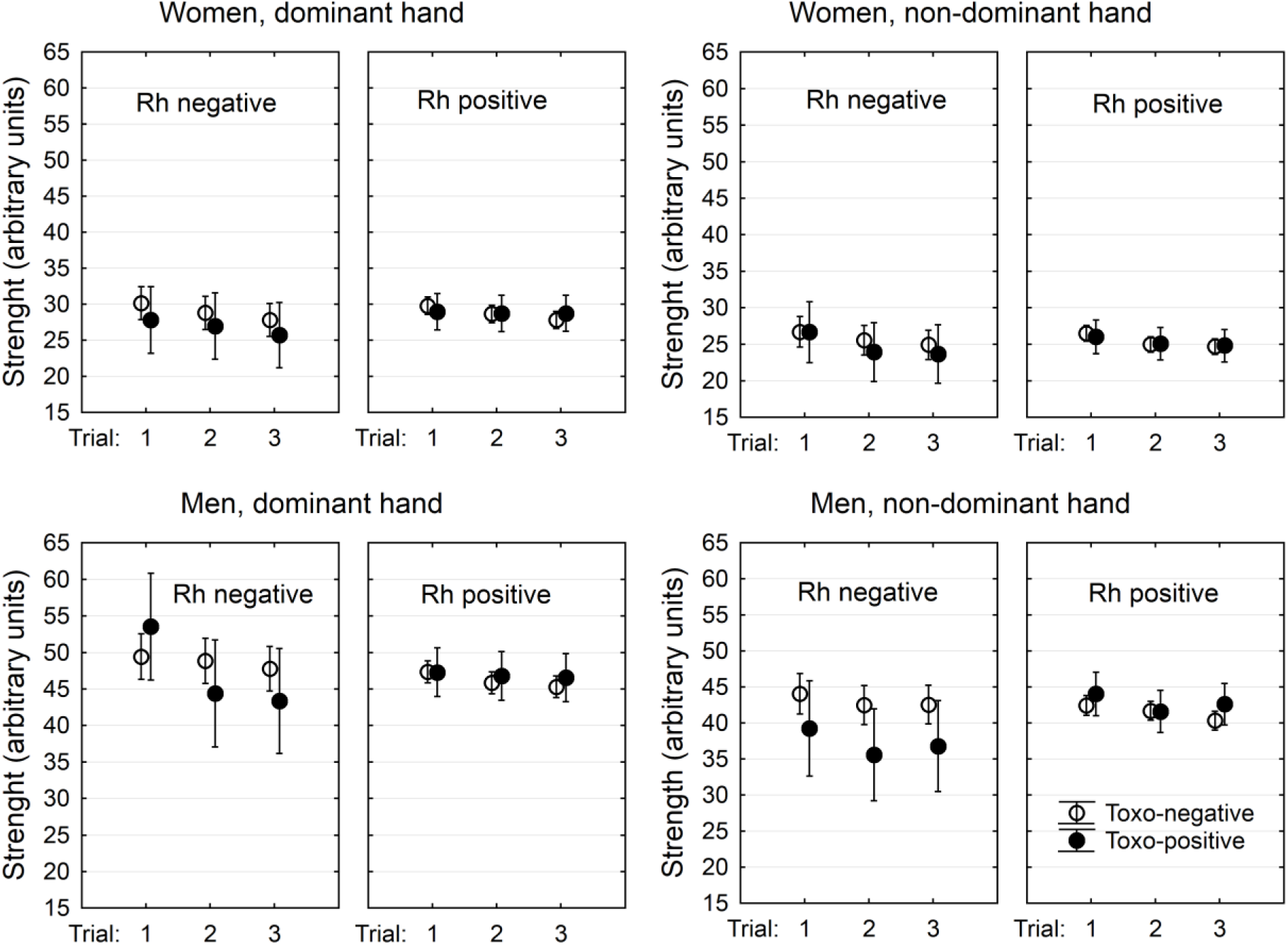
Effects of toxoplasmosis and Rh phenotype on performance in the hand-grip test. Points show arithmetic means of endurance computed for covariate at its mean and spreads denote 95% confidence intervals.

## Discussion

The results of the present study suggest that *Toxoplasma* seropositivity was related to the performance of infected subjects in two tests that primarily measure the isometric (weight-holding test) and isotonic (hand-grip test) strength of participants. This relation was different in Rh-positive and Rh-negative subjects. *Toxoplasma-seropositivity* was always associated negatively with the performance of Rh-negative subjects, but it had a positive association with endurance in the weight holding test of the Rh-positive subjects, especially the men.

The main reason for doing the present study was to find direct evidence for lower endurance of *Toxoplasma-infected* subjects, which has been suspected to exist on the basis of several questionnaire studies (Flegr, 2010). Our results only partly confirmed this prediction as the lower endurance in both tests was observed only in the Rh-negative subjects. Moreover, endurance measured with the weight holding test in the Rh-positive subjects (i.e. in majority of the population) was higher in the infected than in the non-infected subjects. Our experimental setup cannot discriminate between whether the lower performance of the seropositive, Rh-negative subjects in the tests was caused by their lower endurance or by their lower physical strength. However, the fact that better rather than worse performance in the hand-grip test was observed in the first dominant-hand trial in men (see the figure 2) suggests that lower endurance rather than lower physical strength is the more probable explanation of the result.

The existence of significant Rh-Toxo (or Trial-Rh-Toxo) interactions agreed with the results of certain performance tests (Flegr *et al.,* 2008c; Novotná *et al.,* 2008) and of certain health status studies (Flegr *et al.,* 2013; Šebánková & Flegr, 2017) published earlier. We confirmed that *Toxoplasma-infected,* Rh-negative subjects expressed worse results in performance tests, while this was not always true for Rh-positive subjects. In two previous studies performed on blood donors (Novotná *et al.*, 2008) and students (Flegr *et al.*, 2008c), the effects of Rh-toxoplasmosis interaction on performance in simple reaction tests were studied. These studies showed that the *Toxoplasma* free, Rh-negative subjects had shorter reaction times than *Toxoplasma-free* Rh-positive subjects while *Toxoplasma-infected,* Rh-negative subjects had much worse, i.e. longer, reaction times than the *Toxoplasma-infected,* Rh-positive subjects. In the 2008 studies, the authors also showed, but did not discuss, that among the Rh-positive subjects, the *Toxoplasma-infected* ones, especially the Rh-positive heterozygotes, had better performance than the *Toxoplasma-free* subjects. A similar paradoxical phenomenon, namely the positive effect of *Toxoplasma* infection on mental health of the Rh-positive subjects, was described in a case-controls study performed on 79 women who had been tested for toxoplasmosis and Rh phenotype. The *Toxoplasma-infected,* Rh-negative women expressed higher neuroticism measured with the N-70 neuroticism inventory and reported more health problems than the *Toxoplasma-free,* Rh-negative women; the opposite was true for Rh-positive women (Šebánková & Flegr, 2017). It can be speculated that the better performance of the Toxoplasma-seropositive, Rh-positive subjects in performance tests is a result of an increased level of testosterone, which can be associated with higher levels of competitiveness (Archer, 2006). However, while the increased level of testosterone was observed only in men (Flegr *et al.,* 2008a; Flegr *et al.,* 2008b), male mice (Kaňková *et al.,* 2011) and non-castrated male rats (Lim *et al.,* 2013), the better performance of the infected individuals when compared to non-infected individuals in Rh-positive subjects was observed in both men and women. Moreover, the higher competitiveness of *Toxoplasma*-infected subjects can hardly explain the better physical health of infected, Rh-positive subjects (Šebánková & Flegr, 2017). In our evolutionary past in Africa, and probably also in our recent past throughout the world, nearly all people were probably infected with *Toxoplasma.* In Africa 95 % of humans are Rh-positive and in Asia the frequency of Rh-positive phenotypes is even higher, about 99 %. A higher frequency of Rh-negative subjects, about 16 %, is only among Europeans. This was explained by the low abundance of felids and therefore probably the low prevalence of toxoplasmosis in Europe before the recent advent of domestic cat (Flegr *et al.,* 2008c). It is possible that the human body is adapted to this infection and consequently the performance and health of infected Rh-positive subjects is better than the performance of non-infected Rh-positive subjects.

The results of the reaction time and the endurance tests, however, contrast strongly with the results of intelligence tests performed on a cohort of 502 male soldiers (Flegr *et al.,* 2013). This study also showed a significant Rh-toxoplasmosis interaction, however, the infected Rh-negative soldiers scored higher and infected Rh-positive subjects scored lower in two different intelligence tests than corresponding *Toxoplasma-free* subjects. We have no explanation for the results of this study, however, the better performance of the infected Rh-negative subjects in the IQ tests suggest that the effects of Rh-toxoplasmosis interaction on performance are context-dependent and rather specific.

## Limitation

The main limitation of the present study is that the participants have been screened for the Rh phenotype, not Rh genotype. All available data, however, suggest that the Rh-positive heterozygotes, not homozygotes, are resistant to physiological effects of toxoplasmosis. More informative data will probably be obtained when the Rh-positive participants are tested for their Rh genotype. The number of Rh-negative, *Toxoplasma* seropositive male subjects, the rarest subpopulation in general population, is relatively low. This is probably the reason for the absence of a significant effect of the Rh-*Toxoplasma* interaction in males, despite a relatively large effect size. The general design of a study, the cross-sectional cohort study, can prove the existence of statistical association between two or more variables – here *Toxoplasma* seropositivity, Rh phenotype and endurance – but cannot solve the question of causality. Of course, neither *Toxoplasma* seropositivity nor Rh phenotype could be affected by the endurance of a human. However, for example, both the probability of the infection and the endurance can be influenced by some unknown third variable, such as the health status of subjects.

## Conclusions

Our results partly support the hypotheses that latent toxoplasmosis has negative effects on the endurance of infected subjects, and Rh positivity protects people against some negative effects of toxoplasmosis. It must, however, be stressed that all evidence is only indirect and should be supported by the results of manipulative studies performed on laboratory-infected animals.

## Acknowledgements

This work has been supported by Czech Science Fundation 18-13692S and Charles University Research Centre program No. 204056.

